# Predicting the Future with Multi-scale Successor Representations

**DOI:** 10.1101/449470

**Authors:** Ida Momennejad, Marc W. Howard

## Abstract

The successor representation (SR) is a candidate principle for generalization in reinforcement learning, computational accounts of memory, and the structure of neural representations in the hippocampus. Given a sequence of states, the SR learns a predictive representation for every given state that encodes how often, on average, each upcoming state is expected to be visited, even if it is multiple steps ahead. A discount or scale parameter determines how many steps into the future SR’s generalizations reach, enabling rapid value computation, subgoal discovery, and flexible decision-making in large trees. However, SR with a single scale could discard information for predicting both the sequential order of and the distance between states, which are common problems in navigation for animals and artificial agents. Here we propose a solution: an ensemble of SRs with multiple scales. We show that the derivative of multi-scale SR can reconstruct both the sequence of expected future states and estimate distance to goal. This derivative can be computed linearly: we show that a multi-scale SR ensemble is the Laplace transform of future states, and the inverse of this Laplace transform is a biologically plausible linear estimation of the derivative. Multi-scale SR and its derivative could lead to a common principle for how the medial temporal lobe supports both map-based and vector-based navigation.

## Introduction

The reinforcement learning problem is one faced by biological and computational agents alike: finding a series of actions in an environment to maximize long-run reward (Dayan & Balleine, 2002). Many environments have sparse rewards, making a step by step simulation of entire trajectories to choose optimal reward policy (Figure 1, top left) costly and sometimes intractable. One way to make the solution tractable is to generalize representations of nearby states along long paths, allowing simulations to hop over nearby states rather than traverse them one by one. The successor representation offers such a solution (Dayan, 1993). The key idea is that, given a stream of experience and actions, the SR represents a given state in terms of states that will follow it in the near future. Importantly, the definition of “near” depends on a discount parameter, which imposes a timescale on the generalization and hence predictions over successor states (Figure 1, top right). It has been shown that the successor representation (SR) offers a candidate principle for generalization in reinforcement learning (Dayan, 1993; Momennejad, Russek, et al., 2017; Russek, Momennejad, Botvinick, Gershman, & Daw, 2017) and computational accounts of episodic memory and temporal context (Gershman, Moore, Todd, Norman, & Sederberg, 2012), with implications for neural representations in the medial temporal lobe (Stachenfeld, Botvinick, & Gershman, 2017) and the midbrain dopamine system (Gardner, Schoenbaum, & Gershman, 2018).

Most real-world planning problems require agents to plan over longer timescales. Classic reinforcement learning solutions, the model-based and the model-free and hybrid agents, provide non-satisfactory solutions to the problem. A model-free (MF) agent simply stores long-term value of a given action without storing information about the states or the map of the environment. An MF agent solves the RL problem fast, but is inflexible in the face of changes in either the rewards or the map structure of the environment. A typical model-based (MB) solution to finding action policies that maximize rewards is to learn and use a representation of the environment that stores relationships between states that are one-step away from one another, or a one-step transition map of the environment. When the MB agent is about to make a decision, it retrieves different trajectories to reward then computes and compares their expected values via simulating possible sequences one step at a time. Thus, MB agents compute expected value (long-term cumulative reward) for each possible sequence. This one-step MB solution offers flexibility in the face of sudden changes, but it is computationally expensive and sometimes intractable for environments with large state spaces. In contrast, SR offers a more computationally efficient solution to the RL problem by storing generalizations (temporal abstractions) of multi-step relationships between states (Dayan, 1993; Gershman, 2018).

In order to increase efficient computations, SR relies on the weighted sum over the probability of visiting future states, storing expected future occupancy of each future states (that follow from the current state) at a discount scale. This precomputed information about multi-step predictive relationships is later combined with reward to compute value in a linear operation, enabling rapid and flexible adaptation to changes in reward, as in reward revaluation. SR is as flexible as MB when the rewards change in the environment (reward revaluation) but less flexible when the map of the environment changes (transition revaluation), predicting behaviorally asymmetric flexibility. Recent experiments have compared these models to human behavior and shown that the asymmetry in human behavior is more consistent with the predictions of SR agents that update their models via replay (SR-Dyna) (Momennejad, Russek, et al., 2017). It has also been shown that SR offers a computational account of optimal behavior in a variety of RL problems such as policy revaluation and detour (Russek et al., 2017) and explains how context repetition enhances memory-driven predictions in human behavior (Smith, Hasinski, & Sederberg, 2013). SR has also been proposed as a principle for neural organization for place cells and grid cells in the hippocampus and the entorhinal cortex, playing a crucial role in rodent navigation (Stachenfeld et al., 2017). Stachenfeld and colleagues reviewed the literature for the hippocampal-entorhinal encoding of spatial maps during navigation, and modeled the evidence using the successor representation. They concluded that SR is a candidate organizational principles governing the neural firing of place cells and grid cells for learning spatial maps guiding navigation. Since the eigenvectors for the transition matrix and the SR are the same, they suggested that grid cells in the medial entorhinal cortex may provide an eigendecomposition of the graph of the states.

Furthermore, human neuroimaging implicates SR in neural representations underlying event segmentation in the statistical learning of non-spatial relational structures (Schapiro, Rogers, Cordova, Turk-Browne, & Botvinick, 2013). Computational models show that SR can partition the state space, enabling sub-goal processing in large decision trees (Botvinick & Weinstein, 2014). Finally, a recent human fMRI study showed that SR govern the implicit encoding and later retrieval of non-spatial relational knowledge (Garvert, Dolan, & Behrens, 2017). They presented a series of images in an order that was implicitly determined by discrete and non-spatial graphs, and later probed participants about the relation between the different images. They were able to recover the structure of non-spatial relationships from blood oxygen level-dependent adaption in the hippocampus and entorhinal cortex, and showed that the map that best captured behavioral and medial temporal lobe (MTL) results was the weighted sums of future states, as in SR (Garvert et al., 2017).

Taken together, these computational, behavioral, neuroimaging, and electrophysiological evidence in humans and rodents suggest the successor representation as a candidate principle underlying the organization of hippocampal and entorhinal firing in spatial navigation and non-spatial relational learning.

The purpose of our paper is to clarify limitations of existing SR models and propose a solution. Briefly, the limitations are that estimating sequential order and distance between states from an SR with a single discount is nontrivial. Intuitively, for every row of the n×n SR matrix, information about the sequential order and distance between states is lost. SR generalizes over successor states and this temporal abstraction relies on a weighted sum of future states. The weights of successor states exponentially decay the further they are in the future, depending on a discount parameter γ. The parameter γ determines the horizon over which the SR generalizes states. Low γ values discounts states that are further in the future, leading to a shorter temporal horizon over which states are abstracted, while high γ values lead to less steep discounting, and hence a larger time-scale for the abstraction. This weighted sum also depends on the number of times a future state is expected to be visited, starting from the present state.

This can perhaps best be illustrated in an example. Consider the choice between actions that lead from a starting state (e.g., my work) to two different successor states (e.g., two bistros). Depending on how often one or the other is expected to be visited, it is possible that the successor state that is furthest (my favorite bistro) has a higher SR value (expected frequency of future visits) than the successor state nearby (bistro close to work). Depending on the goals of the organism (e.g., how hungry I am and how much time I have for lunch) the optimal choice of a successor state may require separate information about the distances and the frequency of future visits from the present state (e.g., work) to the optimal successor state (e.g., bistro). Thus, due to temporal abstraction, the weighted sum imposes two limitations. The SR vector starting in a given state, and computed with one discount parameter, does not always tell us which future state is closer to the starting location, and what the sequence of states leading to each is. That is, the distance and sequential steps between a starting state and its successor states cannot be computed with merely the SR vector of size. Instead, this estimation requires nonlinear operations involving the entire *n* × *n* SR matrix, involving computation on *n* vectors, each starting in a different state. Given large SR matrices this operation can be costly and it is not likely to be biologically plausible. Our proposed theoretical framework overcomes this limitation by assuming that the brain stores an ensemble of SRs at multiple discounts, and by estimating the derivative of multi-scale SRs.

The simple intuition behind our proposed solution is as follows. Consider an ensemble of *n_g_* matrices of SRs estimated with different *γs*, all between 0 and 1, where *n_g_* is the number of *γ*s and hence SR matrices. Then consider a line going through the jith cell of all SR matrices. This is a vector of expected future visitations from state i to state j across all the SR matrices, with size *n_g_*. Each index in this vector corresponds to temporal abstraction at a different *γ*. The distance between i and state j determines at which index (or *γ*) of this vector the expected future occupancy will change. For an intermediate distance, this relationship could be near zero for low *γ*s but increase for higher *γ*s. For instance, let us say that they are 20 states apart, then their relationship will only register for SRs where the abstraction horizon reaches 20 states into the future, but not for SRs with smaller horizons.

Mathematically, this means that the derivative of this vector of SRs from i to j at multiple discount rates can identify at which distance horizon the relationship between the two states changes, thus recovering their distance. From any given starting state, if we know at which horizon the relationship to every other state changes, it is possible to recover the sequential order and distance of all successor states (see Figure 1). This is a powerful intuition since absent multiple SRs or the derivative, computing distance and order in the bistro example from a single SR requires computing the one-step transition matrix from a given state to all other states using the entire matrix, which is nontrivial.

In short, the derivative of multiple SR matrices can identify at which scales the relationship between two given states change. If we know at which horizon the relationship between every two states i and j changes, it is possible to identify the distance between two states (see Figure 1). In this paper we show that linear operations on an ensemble of multi-scale SRs can estimate their derivatives. Using the derivative, we can reconstruct the sequence of expected future states following a given starting state, and recover order and distance between states, merely by computing a vector as opposed to entire matrices.

In the remainder of the manuscript we offer a more detailed account of our proposal, and show qualitative fit between model predictions and recent neural findings. We conclude with a discussion of the proposal, its neural plausibility, and its possible implications for a unifying account of map-based and vector-based representations using SR.

## An ensemble of successor states is the Laplace transform of the likely future

Where might one look for a gradient of multi-scale SRs, which are equivalent to the Laplace transform? It has been hypothesized that place cells in the hippocampus reflect a predictive map of the environment generated by the SR (Stachenfeld et al., 2017). It is well known that there is a gradient of spatial scales along the long axis of the hippocampus (Jung, Wiener, & McNaughton, 1994; Kjelstrup et al., 2008a; Brunec et al., 2018; Collin, Milivojevic, & Doeller, 2015). For instance, all things equal, CA1 place fields grow in size as the recording location is moved systematically from the dorsal end of the hippocampus to the ventral end (e.g., Kjelstrup et al., 2008a). Furthermore, it has been shown that grid cells are topographically organized in the medial entorhinal cortex (MEC) as well, with the scale of grids increasing from the dorsal border of MEC (Brun et al., 2008). It is possible that this gradient of observed place and gird field sizes corresponds to a gradient of planning horizons over which the SR is computed. Much of the neurophysiology of the MTL hippocampal formation seems oriented to computations along this gradient, suggesting that computation of a derivative is a reasonable computation. For instance, theta oscillations are traveling waves that traverse along the long axis of the hippocampus (Lubenov & Siapas, 2009; Patel, Fujisawa, Berényi, Royer, & Buzsáki, 2012; Zhang & Jacobs, 2015) enabling different spatial scales (and perhaps planning horizons) to be processed sequentially. We discuss the neural plausibility of our proposal in the Discussion section in more detail.

**Figure 1.**
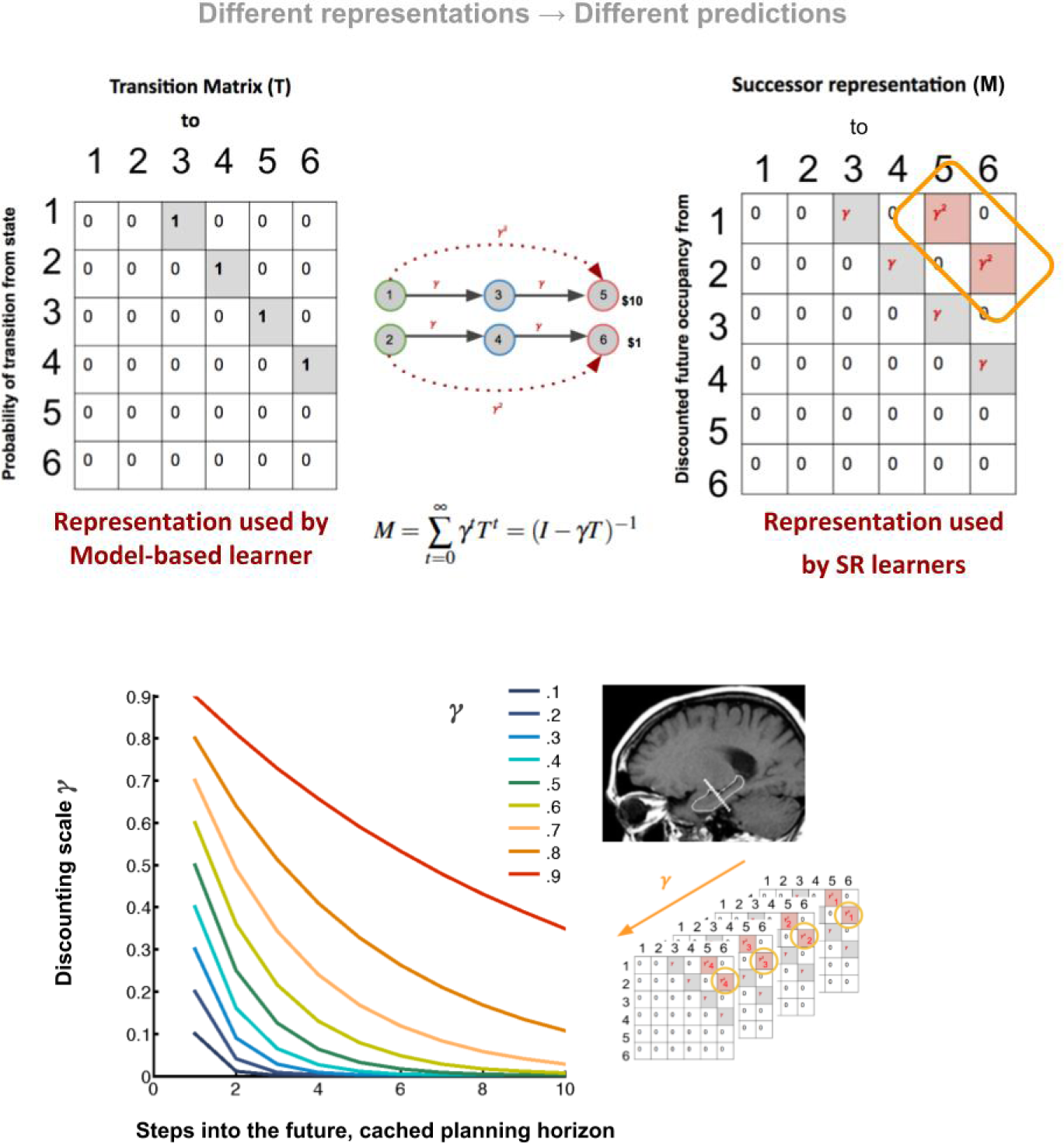
Multi-scale successor representations. (Top) Following Tolman, the classic notion of cognitive maps, and the most common notions of model-based RL, suggest that the brain learns a model of the environment as a 1-step transition matrix. This state-action-state transition matrix (T) stores the probability of transitioning from each state i to state j with a single action a. For instance, it stores the probability 1 for every deterministic transition between the states in the graph of a Markov decision process (MDP) displayed. This 1-step matrix is unrolled at decision time to choose between options, making computation costly, time-consuming, and sometimes intractable for larger decision trees. The successor representation offers a computationally more efficient alternative, where for any starting state i (any row) expected future occupancy of every other state j (each columns) is computed with some discount parameter γ (between 0 and 1). (Bottom) We propose that the brain simultaneously caches multiple successor representations with various scales of discounting. The scales determine the cached planning horizons, can be adaptive to task demands and the volatility of the environment, and can be augmented and updated during wake and sleep offline replay. Neurally, the long-axis of the hippocampus may store the SR ensemble, with smaller scales of abstraction in more posterior regions and higher scales of abstraction gradually spread along the anterior axis. In the following sections we show how such a multi-scale ensemble of SRs can recover both sequential order and distance between states and construct a predicted future trajectory.

In what follows we show that a neurally plausible linear operation, namely the inverse of the Laplace transform, can be used to compute the derivative of multi-scale SR. We first demonstrate that an ensemble of successor representations with different discount rates is the Laplace transform of a timeline of future states. We will then show that a simple inverse of the Laplace transform identifies the derivative of multiple SRs, indicating where the relationships between states change. We show that this operation recovers the sequential order of states and predicts cells that fire at specific distance to goal states, as shown in recent animal and human literature (Sarel, Finkelstein, Las, & Ulanovsky, 2017a; Qasim et al., 2018). We will then discuss the significance of this model for learning and navigation, evidenced by the qualitative match between the model’s predictions and recent findings across species.

The successor representation, described above, provides an exponentially-weighted estimate of future occupancy in going from one state i to state j. That is, when the agent is in its current location in state i, the representation of its successor states are co-activated. The extent of this co-activation depends on the distance: successor states that are nearby are co-activated to a larger extent than those further away, leading to a exponentially-weighted representation of successor states. However, if the multiplied probability of transitioning from state i to a given successor state j is higher than that of transitioning to an equidistant successor state l, then in spite of similar discounting, the state j will have a higher SR value. Therefore, merely on the basis of a row of SR alone, it will not be possible to tell whether state j or state l is closer or not.

It is worth noting that SR is policy dependent, which means that every SR matrix is computed under a given action policy. If the agent expects to visit a state often simply out of habit, the successor representation for that state will have a high magnitude simply due to this policy choice, even if that state does not contain higher reward. This makes it in principle possible that different policies lead to the SR matrix at different time-scales, since the agent may be caching SRs while taking different policies at different scales. This feature may enable multi-scale SR to dynamically select different policy based on changing temporal horizons. In other words, caching a multi-scale ensemble of SRs enables selecting policies with a flexible temporal horizon, which would be more adaptive to task demands compared to a condition where a single predictive horizon was used regardless of the planning horizon required by the task. We return to this point in the discussion.

The choice of discounting and abstraction is controlled by a parameter *γ*, with values between 0 and 1. Choosing different values of *γ* sets the temporal horizon over which predictions can be made. Each successor representation matrix is constructed using a discount rate or scale of generalization, and under a certain policy. Here we propose that the brain constructs multiple SR matrices along the MTL with a broad set of values of *γ* in parallel.

Here we make the point that the choice to use many values of *γ* in parallel also enables a very different form of representation that directly estimates a timeline over future events. This is possible because an ensemble of successor representations with a continuous (or near continuous) spectrum of values of *γ* encodes the Laplace transform of the future. Perhaps an analogy to the better known Fourier transform can provide a good intuition into what a Laplace transform of sequential future trajectories entails. While a Fourier transform decomposes a signal using different sinusoidal functions with different frequencies, the Laplace transform can decompose a signal (e.g., the flow of experience) using exponential curves with different decay rates - note that here we only consider the real part of the Laplace transform.

The Laplace transform has powerful properties, enabling extraction of important information using simple linear operations, such as inversion. The Laplace transform can be inverted with a linear operator 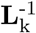 that has been extensively studied in computational cognitive neural models of memory (Shankar & Howard, 2012, 2013; Liu, Tiganj, Hasselmo, & Howard, in press). Importantly, inverting the transform is the equivalent of computing a derivative of the relation between two given states i and j across different SR matrices, i.e., across different time-scales. Intuitively, knowing at which scales the relationship between two states change, indicated by the derivative, enables us to estimate their distance from each other. In short, the multi-scale SR ensemble and its derivative are equivalent, respectively, to the Laplace transform of expected sequential future states and its inversion. Thus, we arrive with cached representation that can also recover the order and distance in estimated future trajectories. To see how this intuition is possible in more detail, we start with a formal description of the successor representation.

Let us suppose that the environment consists of a finite number of states 1..n, or *S*. Depending on the states that an agent visits at every time point, we have a time series of visited states st where each visited state belongs to *S*. The agent visits these states according to a Markov process specified by a one-step transition matrix, where the jith element is the probability of transitioning from state i to state j within one step under action *a*

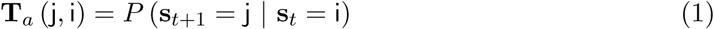

The successor representation (SR) with a given γ value and a given policy, which we denote M (γ), constructs an associative matrix between each pair of states whether they are one step or multiple steps away. The entries in this SR matrix estimate the exponentially-discounted expected future occupancy of any destination state j (any column) when starting at a particular state i, over all future time. That is, the entries of M (γ) estimate:

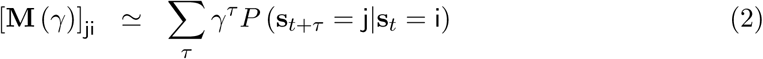

That is,

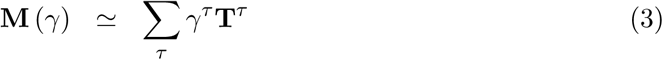

In the case where the statistics of the world are governed by a Markov process, the successor representation can be efficiently computed *via* temporal difference (TD) learning (Dayan, 1993).

Note that Eq. 2 describes a sum over future events. Naturally, the sum contains less information than a complete description of all of the future events. One can readily see this by noting that there are number of possible futures that would give the same value for the sum. For instance, given the same starting state (e.g., work), a visitation to a nearby state that the agent is does not visit often (e.g., nearby bistro) might end up with the same (or even less) weighted SR value as a state with more likely transitions that is many more steps away (e.g., favorite bistro miles away). As such, given that the probabilities of one-step transitions along the trajectory are not all 1 and different policies may be adopted, a higher SR value does not necessarily imply proximity and smaller distance, neither can sequential order be inferred from a comparison of SR values. However, we will show that an ensemble of successor representations with different values of γ can be used to recover the expected future trajectory.

To simplify notation and show equivalence to the Laplace transform, let us define a function that returns the true un-discounted expected trajectory of all future states (whether one or multiple steps away) that are expected to follow a given starting state i, at a lag of time *τ* as:

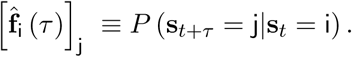

For a successor representation matrix with a single discount rate, M (γ), we can now rewrite Eq. 2 using the function defined above as

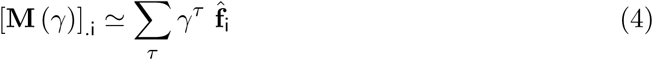

Let us take the continuum limit and define *σ* ≡ − log *γ*. Importantly, *σ* is the real part of the Laplace domain variable. The variable *σ* is understandable as a rate; the time constant 1/σ has units of time and is understandable as the temporal horizon over which predictions are evaluated. Now, we can rewrite Eqs. 3 and 4 as the Laplace transforms of the transition matrix and future following state i respectively:

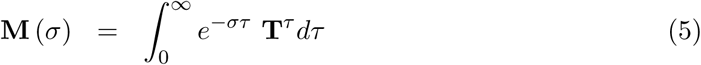

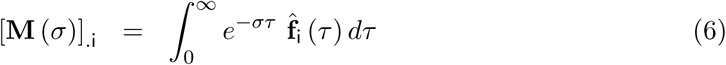

Here we understand the left hand sides of Eq. 6 to refer to many instances of M (*σ*) with a wide range of (real) values of *σ*, i.e., multiple SRs with different (real) values of *γ*. From Eq. 6 it is clear that [M (*σ*)]_i_ is the Laplace transform of the expected states that will follow i as a function of future time, 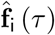.

**Figure 2.**
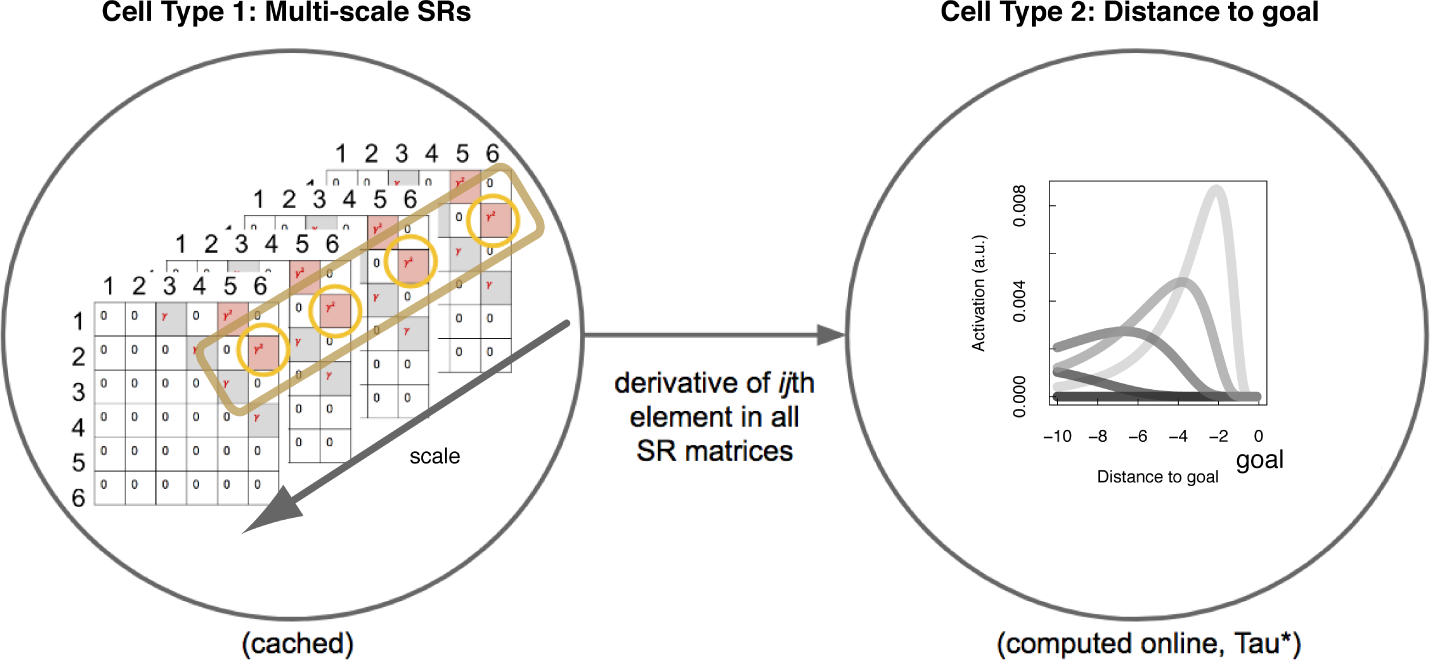
Estimating a timeline of future events from a multi-scale SR ensemble. (Left) The schematic shows an ensemble of SR matrices, each estimated with a different temporal horizon, i.e., a different discount parameter 7. (Right) Computing a derivative as a function of planning horizon for each set of ijth SR value, we construct an activation that rises and then falls as a function of distance from a starting state i to a destination state j. Because different units are responsive to different distances, we can construct an estimate of the distance to each possible outcome. If this is done for all states and all possible distances, across all matrices, we arrive at a series of activations that estimate a future timeline. One method for constructing such an estimate of future timeline is to invert the Laplace transform of the given future timeline, which we show in the manuscript, is equivalent to taking the derivative of the ijth SR ensemble with respect to *σ* ≡ − log γ.

The insight that M (*σ*) is the real Laplace transform of 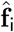 is very powerful. The Laplace transform is invertible; if we could invert the Laplace transform we could explicitly estimate the expected sequential trajectory of future states (Figure 2). The real part of the Laplace transform, which is given by Eq. 6 is sufficient to uniquely specify functions defined over the interval from 0 to ∞. Recall that this sequence of future states was formalized in the future-trajectory function above (see Eq. 4). This means that inverting the Laplace transform could recover an estimate of the function over future events itself.

## Inverting the Laplace transform in a neurally-plausible way

Put another way, Eq. 6 says that the timeline over the sequence of future states 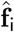 is distributed across different values of *σ*. Our strategy is to invert the transform, recovering this information about the function, and write the answer onto a set of neurons that estimate the future-trajectory function directly. Different neurons in this representation would then estimate the states that are expected to occur at different points in the future. Therefore, if we find a neurally plausible way to invert the Laplace transform, then we will have a powerful tool for recovering a function of future states from a multi-scale but static stack of cached predictive representations.

Fortunately, there has been a great deal of progress on neurally plausible methods to encode and invert the Laplace transform (Shankar & Howard, 2012, 2013). The Post approximation provides a means to estimate the inverse transform using a set of fixed weights that resemble center-surround receptive fields with respect to *σ*. The comparison to receptive fields offers an intuition about how a coarse estimate of the future-trajectory function is arrived at by averaging over a temporal neighborhood in the vicinity of each moment/state along the trajectory.

We denote the set of weights that invert the transform as 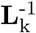, which is defined as

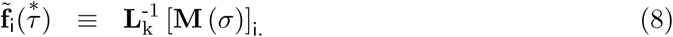

where *C_k_* is a normalization constant that depends on *k*. The *k*th derivative on the right hand side can be approximated numerically. It can be understood in an analogy to center-surround receptive fields. There is a long-standing tradition of identifying center-surround receptive fields using spatial derivatives (e.g., Marr & Hildreth, 1980). In this view, the so-called “Mexican hat” form for the receptive field suggests computation of the second derivative of Gaussian receptive fields.

Putting this together, we construct an estimate of the future following state i by first probing the ensemble M(*σ*) with i. This yields a vector valued (over states) estimate for each value of *σ*. We then operate on this with 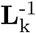, which estimates derivatives with respect to *σ*. The result is a set of units, indexed by 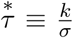. Intuitively, 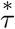 designates the unit of time: the temporal horizon or the horizon of generalization. As we will demonstrate in more detail later, 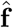, or the future trajectory, is estimated using many units each with a different value of 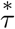:

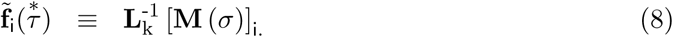

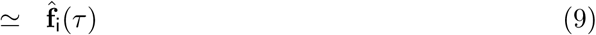

The right-hand of the first line makes reference to the prediction of all future states following from i *via* a successor representation with different values of *σ* [M(*σ*)]._i_. As previously discussed, this is the Laplace transform of the future expected to follow i under a given policy (Eq. 6). This multi-scale successor representation is operated on by 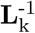, which approximates the inverse Laplace transform (Eq. 7). The second line simply makes the claim that inverting the transform recovers the original function that was transformed, here 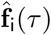. Because 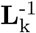 provides an estimate to the inverse transform, this equivalence is only approximate.

The Post approximation ensures that in the limit as *k* → ∞, the approximation of future states becomes perfect (Post, 1930). When *k* is finite, 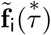 is a coarse-grained estimate of 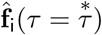 averaging over a temporal neighborhood in the vicinity of *τ*. Following (Shankar & Howard, 2012) it can be shown that the width of this coarse-graining is exactly proportional to each units value of 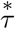, implementing a scale-invariant estimate of the future.

## Simulating points along a future trajectory

The form of the curves representing the future can be readily understood analytically starting from the Laplace transform of a delta function. Consider the case where some state perfectly predicts an outcome some time *τ*_0_ in the future. This would obtain if a set of states are deterministically experienced in a long sequence. Let us write 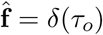 to describe the function describing the distance to the goal state from the present. In this case, we find the Laplace transform of this function, generated by the successor representation is given by

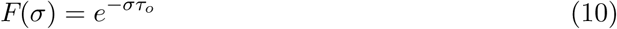

The left panel of Figure 3 shows this expression for different values of *σ* (different curves) with the distance to the goal state *τ_o_* on the *x*-axis. This describes the activation of these units unfolding as the agent comes closer to the goal state moving left to right.

Now, to compute 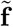 we must take derivatives of *F*(*σ*) with respect to *σ*. Each derivative of Eq. 10 spits out another factor of *τ_o_* and we find (including the remainder of the definition in Eq. 7),

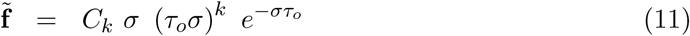

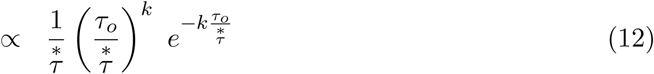

where we have used the definition of 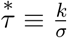 in the second line. The right panel of Figure 3 shows this expression for units with different values of 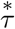 (different curves each corresponding to a different horizon unit) as the goal state is approached. Rather than increasing monotonically as the goal is approached, neurons in 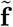 peak in their activation when the goal is a characteristic distance in the future.

The form of Equation 12 makes several properties of the representation clear. The expression is the product of two terms, an increasing power law function 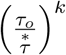 and a decreasing exponential term 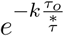. The product is zero when the ratio is zero and also when the ratio is infinity. As *k* increases each of these terms becomes more steep. An elementary calculation shows that Eq. 12 is maximal at 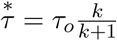. Considering the behavior of Eq. 12 as one varies *τ_o_*, the unit coding for 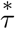 steps in the future is maximally activated when 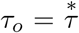, independent of *k*. Because τ_o_ appears only as a ratio with 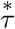, we can see that the model is scale-invariant. That is, starting for any particular value of *τ_o_* (and *k*), we find some pattern of activation as a function of 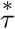. If we change the value of *τ_o_* to 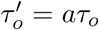, we will find the same pattern of activation across units (up to a constant factor) by remapping 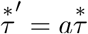.

The goal of Figure 4 is to provide mechanistic insight into why 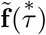 has the properties it does. Rather than showing the activation of units unfolding in time Figure 4 shows the activation across all units in the multi-scale SR (*x*-axis) and 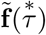 at different moments when the goal state is predicted at different distances in the future (different lines). The top left panel shows the multi-scale SR for each possible value of γ. In each case, one obtains an exponential function; when the goal is closer, the curve is shallow (lighter lines); when it is further in the future the curve is more steep (darker lines). The curvature of this function over values of *γ* encodes information about when the outcome is expected to occur. In order to decode this information in units of distance to the goal, we need to change variables from the discount factor to units of distance. *σ* is understandable as a rate; *k/σ* has the same units as τ*_o_*. The bottom left shows the mapping between *γ* and *k*/*σ*. The y-axis of this graph is truncated. This is because as *γ* goes to 1 the “time constant” *k/σ* grows without bound corresponding to an infinitely broad planning horizon.

**Figure 3.**
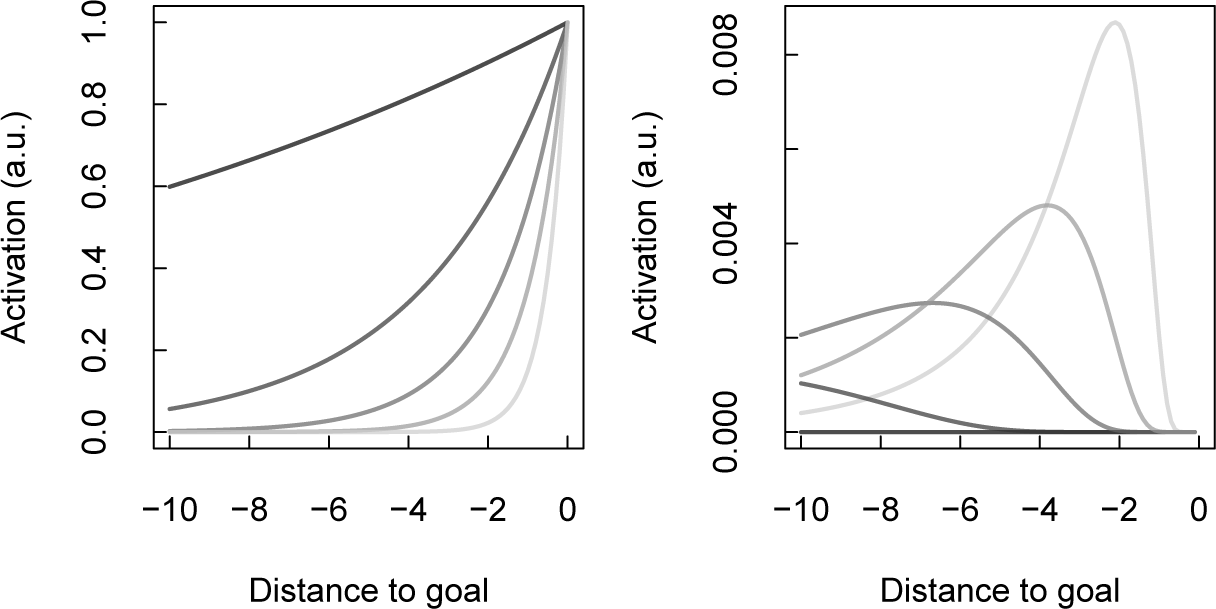
Activation of neurons representing the goal state in the multi-scale SR (left) and in the estimate of future timeline (right). The panels correspond to activations in two different types of neurons. In each panel, the activations of units or neurons coding for a goal state are shown as the agent approaches the goal along a 1-dimensional sequence of states. The x-axis shows the distance to the goal. The y-axis shows goal activation as the path is traversed. In the left panel, different shades of the curves correspond to different values of γ and on the right, they correspond to different values of 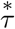 or different horizon unit (right). Higher values of γ are shown in darker shades of grey (left). Higher values of 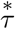 are shown in darker shades of grey (right). As can be seen by the shape of the curves on the left, units in the multi-scale SR show an exponential increase as the agent approaches the goal; larger values of *γ* result in a less sharp increase. Units in the future timeline (right) also predict the goal state and change their activation as the goal is predicted at different distances in the future. However, rather than increasing monotonically, these units peak when the goal is predicted in a particular future horizon, i.e., a particular number of steps or distance in the future. Neurons with larger values of 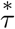 (darker curves) peak when the goal is further away in time. Neurons with smaller values of 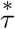 peak when the goal is closer.

The upper right plot in Figure 4 replots the data from the upper left curve, but as a function of *k/σ* rather than γ (the axes are also rescaled). Note that each of these curves start at zero with a very shallow slope. Moving to the right, they shift to a regime with a non-zero slope. Note that the second derivative of each curve would be very small at the left of the plot and very small at the right of the plot. In between, the curve rapidly changes from the shallow slope to the higher slope. The inflection point where this happens depends on how far in the future the goal state is predicted. Because each unit in 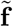 indexed by 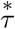 computes derivatives with respect to *σ* in the neighborhood of 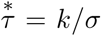, these units are maximally activated when the goal state is expected 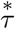 units in the future. With many values of 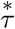, the pattern of 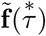 over different values provides a veridical but coarse-grained estimate of when the goal state will be observed in the future subject to the policy that generated the multi-scale SR.

## Predictive maps and distance to goal firing patterns

Multi-scale successor representations can be thought of as a set of predictive maps, each with a different scale or predictive horizon. The analytic expression above (Eq. 12) can be readily calculated for simple environments for which the multi-scale SR has a simple closed form solution. However, operating on the multi-scale SR with 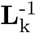 constructs an estimate of expected future outcomes as a function of distance across expected paths given a particular policy. Figure 6 illustrates the multi-scale SR and the estimate of future timeline for a gridworld environment with a barrier.

**Figure 4.**
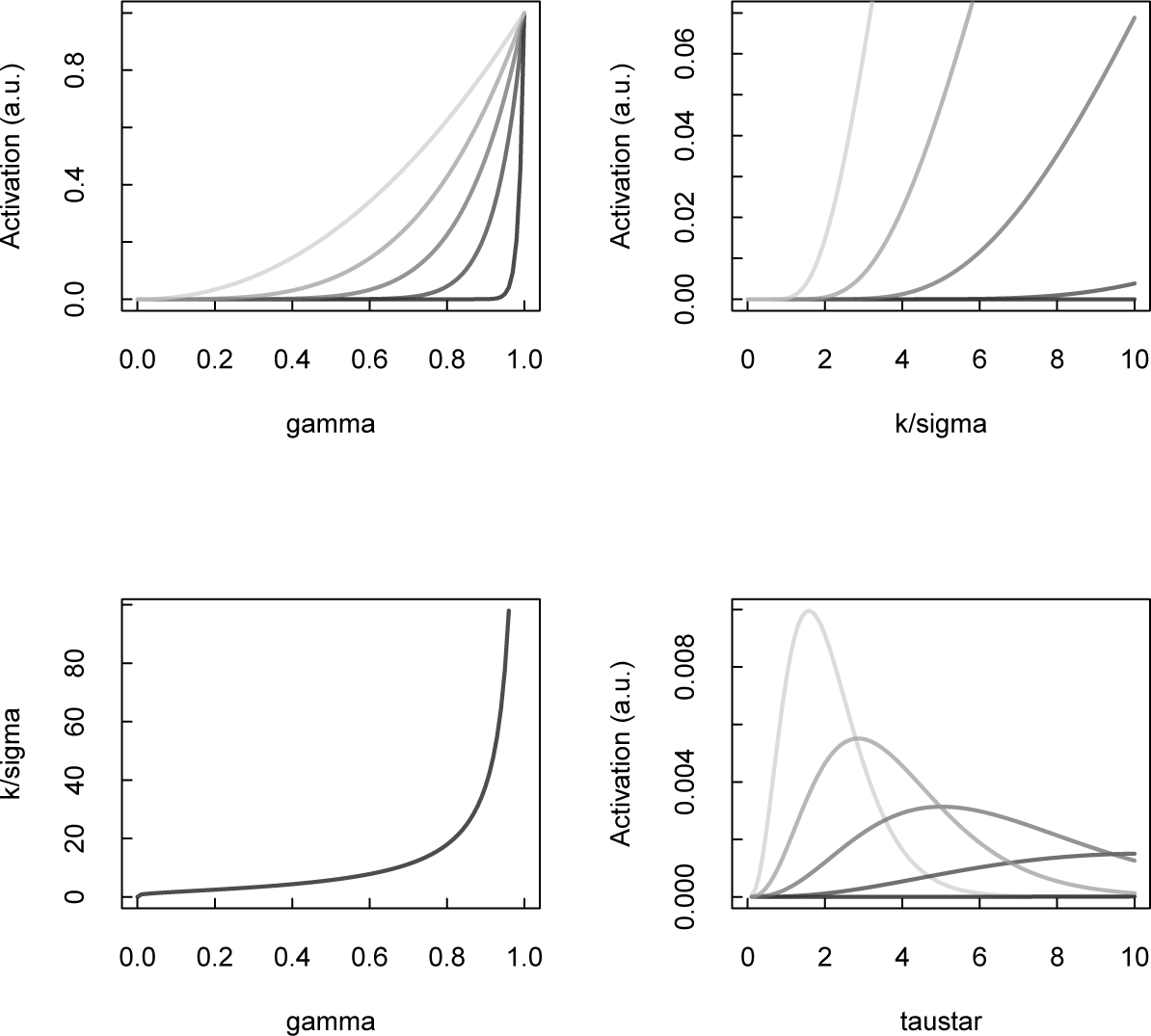
Top left: Prediction of the goal state by the multi-scale SR as a function of γ when the goal state is different distances in the future (separate curves). Darker curves correspond to cases when the goal is further away. In each case the multi-scale SR gives rise to a gradient over different values of γ; the distance to the goal is encoded by the steepness of this curve. Bottom left: The discount factor γ is in a one-to-one relationship with the Laplace domain variable *σ*. The curve in the bottom left shows *k/σ* as a function of γ. Note that *k/σ* grows without bound as γ goes to 1. Top right: The curve in the top left is shown as a function of *k/σ* rather than as a function of γ. The y-axis is compressed to better show the detail of the activation. The x-axis is chosen to go up to 10. Note that the curves start at zero and then rise around an inflection point. The position of the inflection point depends on the distance to the goal. When the goal is closer (lighter curves), the inflection point is found at smaller values of *k/σ*. Bottom right: The estimate of future outcomes as a function of 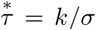 is constructed by taking the derivative with respect to *σ*. he axes of the right panels are chosen so that the units align with one another. The estimate of the future is computed by taking derivatives with respect to *σ* in the neighborhood of the corresponding value of *k/σ* (see Eq. 7). Because the inflection point in the top right panel depends on the distance to the goal, computing derivatives with respect to *σ* results in a representation of the distance to the goal as function of the future.

To construct the SR, one first starts with the transition matrix T which is non-zero for each transition between adjacent points that is not prevented by a barrier. This can be a policy-independent or policy-dependent transition matrix. As an agent navigates the environment, SRs should be estimated based on the agent’s experience and in a policy-dependent manner, where successor states are dependent on the agent’s action policies. For the present purposes, however, we will use an equation that obtains for a random-walk version of SR. For a random policy, it is possible to estimate SR simply from a one-step transition matrix using the equation below. Computing SRs using the one-step transition matrix is the equivalent of computing SR while the agent navigates the environment under a random policy, if the rewards were equally distributed across the environment. Following previous work (e.g., Dayan, 1993; Momennejad, Russek, et al., 2017; Stachenfeld et al., 2017), here we computed successor representations at multiple scales from the one-step transition probability matrix (**T**) according to

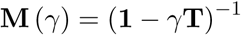

Figure 5 displays a multi-scale ensemble of SRs with different γs as well as the corresponding derivatives (inverse of transforms for various values of *σ*) for a gridworld with a π-shaped barrier in the middle, and reward in position (2, 2). The plots on the left show an ensemble of predictive maps, i.e., M(γ), predicting the successors of a state in the upper left of the gridworld with different values of γ. As γ increases, the gradient across states becomes more shallow; at large horizons one can note the π shaped barrier. We applied 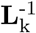, computed with a discrete approximation to the derivative operator (Shankar & Howard, 2013), to these predictive maps and obtained the panels on the right of Figure 5. States of the same shade in these panels represent states that are the same expected distance from the goal state subject to the policy used to generate the **M**(γ). This procedure naturally takes into account barriers—note that one can read off that states close to the goal but on the other side of the barrier are the same distance in terms of expected paths as states in the lower right corner of the environment.

There is some evidence that these computational properties are reflected in the firing patterns of cells in the hippocampal and medial temporal formations in animal models. As shown above, computing the inverse (or the derivative) of the multi-scale SR ensemble resembles firing patterns at specific distances to any given destination state. This predicts sequential activations of different units/cells as a function of distance to the goal (this ‘goal’ could be a goal object, a reward state, a frequently visited location or destination, a specific boundary, a subgoal, a remembered location, etc.). Whereas the successor representation itself gives monotonically decreasing gradients into the future (like border cells), the inverse of a multi-scale SR ensemble predicts cells that are sequentially activated as a function of distance to goal (these can be thought of as distance-to-goal cells).

These theoretical predictions offer a qualitative fit (though we did not compute a quantitative fit) to three recent empirical findings in the bat, rodents, and humans. These studies have discovered cells that display sequential firing as a function of distance to a goal state. First, Sarel and colleagues (Sarel, Finkelstein, Las, & Ulanovsky, 2017b) investigated the representation of spatial goals in the hippocampus of bats (Figure 6, right). They reported that CA1 neurons of flying bats displayed angular tuning to the goal direction, many of which were tuned to the goal even when it was occluded. Importantly, some of these cells encoded distance to goal as well as goal direction. The distance to goal aspect is in line with earlier reports of goal-approach cells in rodents (Eichenbaum, Kuperstein, Fagan, & Nagode, 1987). These firing patterns are consistent with the idea of a vectorial representation of goals in the hippocampus. Our simulation of distance to goal firing based on our theoretical proposal in Figure 6 resembles the distance to goal cells in the bats.

**Figure 5.**
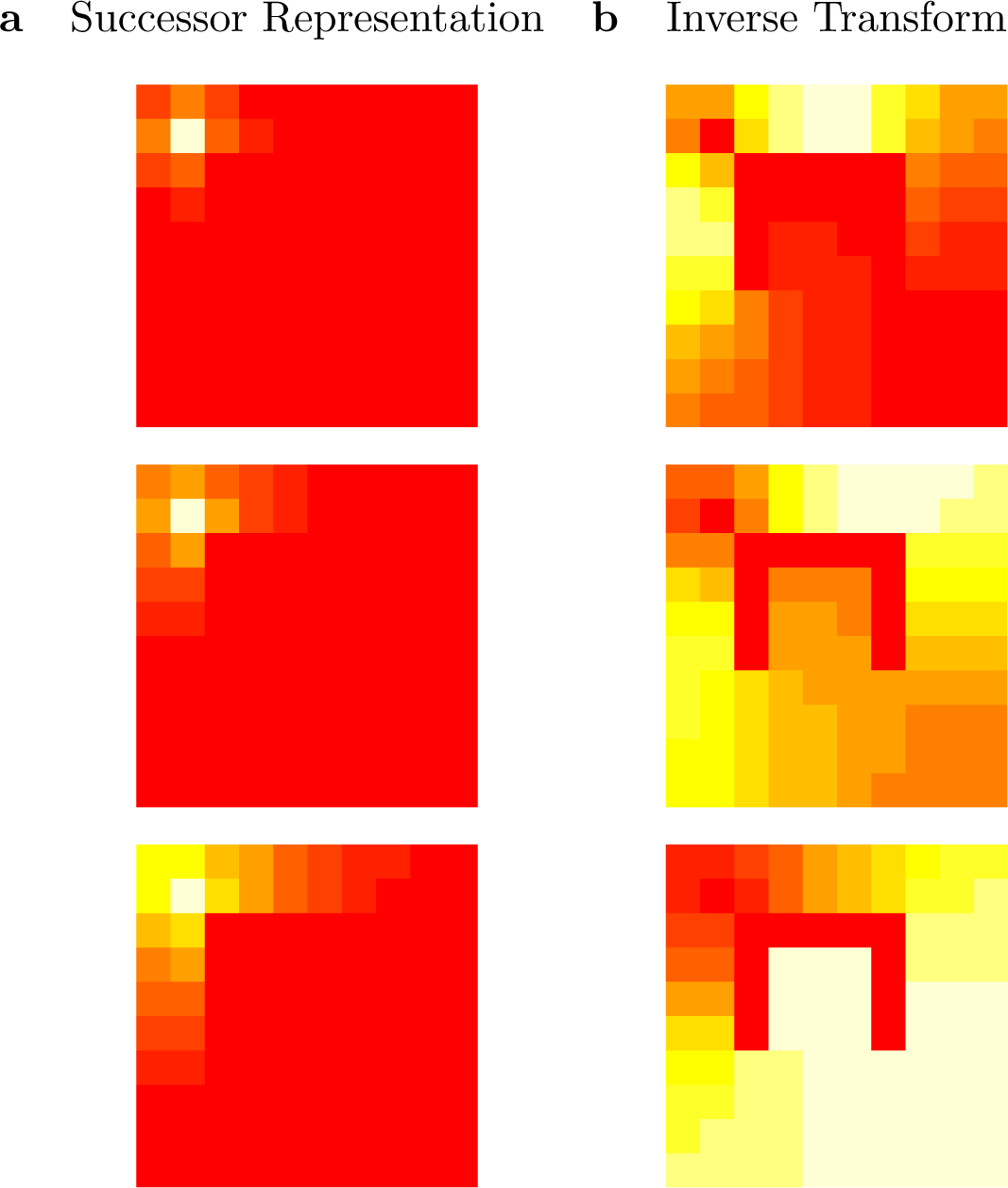
The successor representation and inverse transform in a 2-D gridworld with a *π*. shaped barrier near the center. The axes correspond to x and y coordinates uniquely identifying each cell in the grid world. A reward is placed on the top left cell corresponding to location 2, 2. Each image shows the value of the successor representation starting from a location near the upper left of a gridworld to every other cell in the grid world. Higher SR values, or higher expectations of future occupancy, are reflected as brighter colors. **a**. Successor representation matrices for three different values of γ. The brightest location in the top left is the location of the reward, or goal state. As γ increases (top to bottom), there is markedly higher expected occupancy for further states that lead to the goal. For all values of γ, future occupancy decays monotonically from the starting point along possible paths. SR matrices were normalized to their peak value. **b**. Inverse transform for corresponding values of 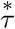 each of which corresponds to a temporal field during the process of inverting the SR ensemble. To obtain the inverted Laplace transform in this figure we set *k* = 8. Note that rather than decreasing monotonically, successively larger values of 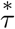 (top to bottom) lead to activation in positions within the gridworld with further distances to the goal state.

**Figure 6.**
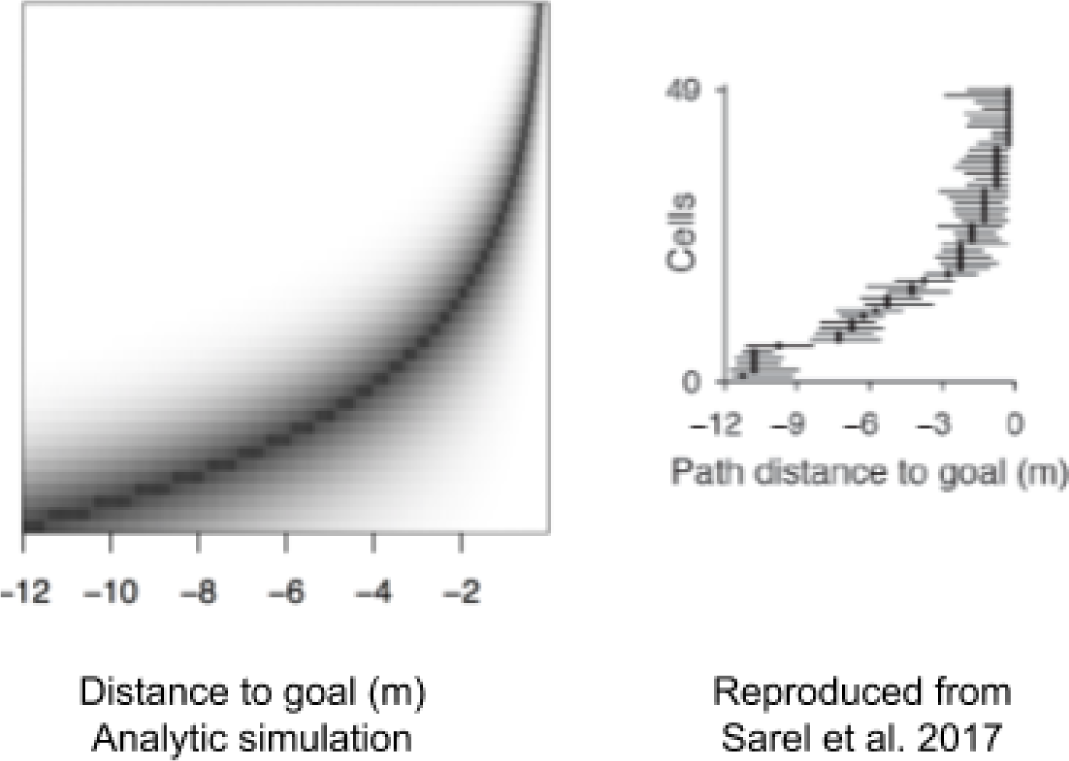
Qualitative comparison of units with distance-to-goal firing in the future timeline (Eq. 12) and distance-to-goal cells in the bat hippocampus. (Left) Constructing a future timeline from an ensemble of multi-scale SR maps predicts units that each fire within a different distance to the goal state. The y axis here corresponds to different units, and the x axis to the simulated distance from the grid world shown in the previous figure above. (Right) Firing patterns of goal-cells discovered in the bat hippocampus, reproduced from Sarel et al., 2017. Note the resemblance to the predicted distance-to-goal firing on the left.

The second piece of evidence consistent with distance-to-goal firing comes from Gauthier and Tank (2018), who report mouse CA1 cells that tune to specific distances from reward locations (which they refer to as reward cells). The study reports that these reward cells were tuned to similar distances to reward states even when the animal was moved to another environment. This is what one would expect from this conception of a future timeline. Because the same outcome is predicted at the same distance in the two environments, this should result in the same cells being activated to code for the same future outcome.

The third piece of evidence comes from human electrophysiology (Qasim et al., 2018). Qasim and colleagues report cells in the human entorhinal cortex (EC) that fire at certain distances as people approach remembered locations during cued object-location recall. In this case, the goal location is not visible in the environment, and is instead an internally generated goal related to the memory retrieval cue.

In short, consistent with our predicted distance-to-goal firing cells, recent empirical evidence suggests cells that fire at specific distances to goal in the hippocampus and entorhinal cortex of bats (Sarel et al., 2017b), rodents (Aronov, Nevers, & Tank, 2017; Gauthier & Tank, 2018), and humans (Qasim et al., 2018). As indicated in Figure 6 the inverse (or derivative with respect to *σ*) of a multi-scale SR ensemble simulates the reported distance to goal effect. One difference between the bat goal-cell findings and our distance-to-goal units is that, in the bat results, more cells fire in closer proximity to the goal state (see Figure 6, right). Furthermore, in the model the units that fire in closer distance to the goal states show less uncertainty. This could be because our model overestimates certainty about what state the agent is in. This can be addressed by including a certainty term. Taken together, this correspondence suggests that vectorial representations could emerge from static cached maps. (Johnson & Redish, 2007; Pfeiffer & Foster, 2013).

Our multi-scale successor representation framework could potentially offer a unified principle that supports both the map-like representations elicited by place cells, asymmetric firing skewed towards the goal location (Stachenfeld et al., 2017), and the vectorial representation of direction and distance to goal (Kubie & Fenton, 2009; Bush, Barry, Manson, & Burgess, 2015). The predictions of this proposal are in line with the observation of distance to goal cells (Fig. 6).

## Discussion

The successor representation (SR) offers a principle for abstract organization in reinforcement learning (Dayan, 1993; Momennejad, Russek, et al., 2017; Russek et al., 2017), computational accounts of episodic memory and temporal context (Gershman et al., 2012), and predictive representations in place cells and grid cells in the hippocampus and medial entorhinal cortex (Stachenfeld et al., 2017). The successor representation (SR) offers a solution to planning at large temporal horizons and optimal sub-goal discovery (Botvinick & Weinstein, 2014). Combined with offline replay, a model known as SR-Dyna is superior to classic model-free and model-based reinforcement learning mechanisms in explaining human behavior (Momennejad, Russek, et al., 2017), outperforming hybrid MF-MB models and varieties of earlier Dyna models across other problems as well (Russek et al., 2017). Neurally, it can also explain asymmetric firing toward the goal state in hippocampal place cells (Stachenfeld et al., 2017). However, on its own, a single row of SR discards fine grained information about order and distance of a starting state to expected future states. This information is important to animals in many real-world decision making problems, and call for an adequate account in any computational proposal.

Here we have shown that a multi-scale ensemble of successor representations can overcome these limitations: the derivative of the SR ensemble can estimate the expected sequence of future states following a starting state, recovering both sequential order and distance between states. Importantly, this derivative marks changes in the relationship between two given states across the timescales of abstraction. How can the brain compute this derivative in a neurally plausible manner? We have shown that a multi-scale SR ensemble is equivalent to the real Laplace transform of a given states timeline of successor states. The inverse of this Laplace transform computes the derivative of the SR ensemble, recovering which future states lie within given temporal horizons of a given state (e.g., the present state, or the goal state). Importantly, the Laplace formulation and its inverse are closely related to an established neurally plausible proposal for a scale-invariant representation of the past in the medial temporal lobe (Howard et al., 2014). In short, we mathematically show the neural plausibility of the idea that the brain may store multi-scale SRs and compute their derivative, leading to both predictive map representations and neural firings akin to vectorial representation.

Notably, the inverse of the Laplace transform (i.e., derivative of the multi-scale SR) predicts sequential firing of “distance-to-goal” neurons when the agent is in certain temporal neighborhoods of (or distance to) the goal state (Figures 3 and 6). We show that this analytic model prediction resembles recent findings in bats, rodents, and humans (Sarel et al., 2017b; Gauthier & Tank, 2018; Qasim et al., 2018) that are consistent with the idea of vectorial navigation. The rodent data calls these cells reward cells, the bat data has been taken these cells as evidence for goal-vector cells, and the human stury refers to them as trace cells. What these results and our proposal here have in common are units/cells that each fire at a specific predictive distance to a goal state, such as reward in the rodent study, resting locations (or destination state) in the bat study, or remembered goal locations in the human study. What our proposal further provides is that, computationally, all these varieties of vectorial representations supervene on underlying predictive maps. Thus, our proposal combines vector-based and map based navigation in a neurally plausible fashion.

### Neural plausibility: Evidence from place cells, grid cells, and time cells

We have suggested that the derivative of a multi-scale SR ensemble offers a linear solution for computing the distance and sequential order of a given state’s future trajectory. But is multi-scale SR neurally plausible? The proposal that SR can be stored at multiple scales along the long axis of the hippocampus has been suggested elsewhere as well (Stachenfeld et al., 2017) and is in line with the following observations in the rodent literature. First, it has been shown that hippocampal place cells, i.e., neurons the activity of which corresponds to the animal’s current location, fire over a larger spatial radius in more ventral (anterior in humans) hippocampal neurons compared to more dorsal (posterior in humans) hippocampal neurons. In other words, the size of place fields vary along the long axis of the hippocampus, with topologically more anterior hippocampal regions representing larger place fields (Kjelstrup et al., 2008b). Furthermore, grid cells are topographically organized in the medial temporal lobe (MTL) as well, and the scale of the grids has been shown to increase along the dorsal border of the medial entorhinal cortex (Brun et al., 2008).

It has also been shown that the so called place cells also respond to non-spatial relations, e.g., sonic and social state-spaces (Aronov et al., 2017; Omer, Maimon, Las, & Ulanovsky, 2018). This suggests a potentially non-spatial underlying computational role for place and grid systems that is not restricted to spatial navigation but can support the organization of non-spatial relational knowledge as well (Garvert et al., 2017). Based on these findings it is possible that the brain learns an ensemble of successor representations along the long axis of the hippocampus, each with a different discount parameter corresponding to different scales of abstraction, providing information about progressively larger horizons along the long axis of the hippocampus (see Figure 1). Notably, topologically organized place fields have also been observed in cortical regions such as the retrospenial cortex, but it has been shown that these fields develop slowly over time and are hippocampally dependent: they are attenuated with hippocampal damage (Mao et al., 2018). Similarly, while multi-scale SRs can also be encoded or consolidated in the neocortex, it is likely that the hippocampus plays a crucial role in their formation.

In addition to encoding spatial representations, MTL neurons also carry explicit information about the time at which events were experienced in the past. Neurons referred to as time cells have been observed in many brain regions (Pastalkova, Itskov, Amarasingham, & Buzsaki, 2008; Jin, Fujii, & Graybiel, 2009; MacDonald, Lepage, Eden, & Eichenbaum, 2011; Kraus et al., 2015; Mello, Soares, & Paton, 2015; Akhlaghpour et al., 2016; Tiganj, Kim, Jung, & Howard, 2017; Tiganj, Cromer, Roy, Miller, & Howard, 2018). These neurons appear to show receptive fields in time for past events; as a stimulus recedes into the past different neurons become sequentially activated, presumably as the stimulus sequentially enters their receptive fields. Because each neuron has a temporally circumscribed “time field”, enabling the animal to learn associations between stimuli separated in time via a simple association. Looking across neurons, it is possible to directly read off an estimate of the time at which different events were experienced. Time cells may rely on representations of events that decay at different rates, allowing an estimate of past events that extend backward from the present moment. If one could construct an analogous estimate of future events that extend forward from the present, this would provide us with explicit sequential information about future events. Such an estimate would resemble distance to goal cells, which fire at particular distances to a goal location (Eichenbaum et al., 1987; Sarel, Finkelstein, Las, & Ulanovsky, 2017c; Gauthier & Tank, 2018; Qasim et al., 2018). As shown above, the derivatives of multi-scale successor representations may offer a theoretical account for the emergence of such firing from predictive maps.

While sufficient neurophysiological data is not available to strongly constrain our account, progress in understanding the neurobiology of the Laplace transform for memory of the past may provide some insight. Shankar and Howard (2012, 2013) argued that the brain could construct a memory as a function of how far in the past events were experienced *via* the Laplace transform. Analogous to the present paper, a set of cells code the Laplace transform of the past. These cells project *via* 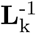 to another set of cells that approximate functions of past time.

Neurons coding for the time of past events would behave like so-called “time cells” that had been observed in the medial temporal lobe (Pastalkova et al., 2008; MacDonald et al., 2011). Since the initial reports, time cells have been observed in a wide range of regions, tasks, and species, confirming several qualitative predictions of the theoretical account via the inverse Laplace transform (e.g., Tiganj et al., 2017; Tiganj, Cromer, et al., 2018; Mello et al., 2015; Akhlaghpour et al., 2016). Of course, time cells that align with the properties predicted for the inverse Laplace transform of the past could have been constructed via some other mechanism. However, recent preliminary work shows evidence for neurons that decay exponentially with a spectrum of rates in the entorhinal cortex (Tsao et al., 2017; Meister & Buffalo, 2017); just the form predicted for the Laplace transform of time. Notably, “the Laplace transform of the past” could be understood in the context of reinforcement learning as “a set of eligibility traces with different decay rates.”

Given the neural plausibility of the Laplace transform itself, the only other requirement to neurally construct the inverse is to find a biological basis for the operator 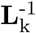. This operator can be understood as a set of receptive fields conveying information from cells encoding the transform (indexed by *σ*) to cells estimating the inverse (indexed by 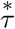). It has long been understood that center-surround receptive fields compute derivatives (e.g., Marr & Hildreth, 1980) much like those required for eq. 7). The entries in 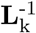 can be understood as describing center-surround receptive fields with respect to *σ* (Liu et al., in press). In the visual system it has been understood that general computational principles could enable learning of center-surround receptive fields from natural statistics (Bell & Sejnowski, 1997; Olshausen & Field, 1996). These findings suggest that the the operator 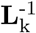 used here is neurally plausible.

### The distribution of scales and discretization of time

What have attributed time-scales to the parameter *γ*. It is not obvious whether the distribution of *γ* scales is approximately continuous or discrete. The mammalian brain provides both discrete and continuous distributions of scales for related quantities. For instance, the distribution of spatial frequency of grid cells appears to be organized into discrete modules along the entorhinal cortex (Barry, Hayman, Burgess, & Jeffery, 2007; Stachenfeld et al., 2017). These modules consist of many grid cells with the same spatial frequency but different spatial phases. The modules are organized along the dorso-ventral axis. In contrast, the distribution of time cell receptive fields appears to be continuous even within dorsal CA1 (Mau et al., 2018). That is, within the population of time cells in CA1, the set of time cells continuously maps the delay interval (up to at least a few dozen seconds). On another note, a recent study has shown that temporal information is robustly encoded across time scales (seconds to hours) in lateral entorhinal cortex populations (Tsao et al., 2018), while similar information was not found in either CA3-CA1 nor in the medial entorhinal cortex. It is, however, computationally possible that a small number of discrete nodes over future time provides sufficient information to solve computational problems related to time-scales. For instance, a deep Q-learning network performing a video game task was able to learn efficiently with just a handful of logarithmically-sampled points of a temporal history (Spears, Jacques, Howard, & Sederberg, 2017). Future work is required to shed light on the distribution of scales and the discretization of time.

### Relationship to prior modeling work

We have proposed that the brain maintains successor representations at a spectrum of scales or *γ*s. Previous modeling works have proposed spectral assumptions about learning rates and eligibility traces. In (Kurth-Nelson & Redish, 2009), the authors used a set of “micro-agents” each with a different learning rate. They showed that the set of micro-agents, with a distribution of learning rates across agents, was able to simulate behavior akin to hyperbolic discounting. Ludvig, Sutton, and Kehoe (2008) propose a model in which the decaying trace of a stimulus is approximated by a series of basis functions with receptive fields spread across trace heights. The time course of the trace results in “microstimuli” that get shorter and wider with time. Future modeling work can better illuminate the relationship between spectral proposals about learning rates, eligibility traces, and the scale parameter in SR.

Other recent models have proposed methods for constructing a compressed estimate of future events by exploiting properties of the Laplace transform. Tiganj, Gershman, Sederberg, and Howard (2018) constructed an estimate of future events as a function of time using a simple associative account in which a compressed estimate of the past—analogous to a set of sequentially-activated hippocampal time cells—was associated to the present. The associative operator learns simple Hebbian associations between the past at each lag 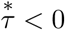 and the present. In order to estimate the future a lag 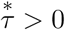 in the future, the present stimulus is used to probe the appropriate association. Tiganj, Gershman, et al. (2018) described a number of computational properties of this method, including scale-invariance. Shankar, Singh, and Howard (2016) exploited the properties of the Laplace domain to translate the current estimate of the past to estimate the future. That model made a detailed mapping between function translation and theta phase precession. Although the endpoint of these models—a scale-invariant estimate of future events that makes order and distance explicit—is quite similar to the endpoint of the method developed in this paper, the mechanisms used to generate them are quite different. The multiscale SR described here can use efficient temporal-difference learning algorithms to construct the SRs. In environments where the world is well-approximated by a Markov process, this property can represent a considerable computational advantage over associative models.

### Relationship between map-based and vector-based navigation

An interesting implication of our account is that it supports properties of both map-based and vector-based navigation. Map-based navigation relies on computations using an underlying cognitive map like representations of states, which in our proposal take the form of multi-scale predictive representations. Vector-based navigation, on the other hand, enables rapid planning of direct trajectories to goals via estimates of distance and direction to the goal state (Kubie & Fenton, 2009; Bush et al., 2015). Previous work has contrasted map-based navigation with vector-based navigation for solving optimal route to goal in path integration problems (Kubie & Fenton, 2009). It has been proposed that vector-based navigation requires computing a “shortcut matrix” in memory: a set of shortcut vectors computed for pairs of visited locations (Kubie & Fenton, 2009). Kubie and Fenton contrast insect navigation, where heading vectors are computed by path integration, with mammalian navigation where hippocampal place cells are typically considered to govern map-based navigation. However, when considering map-based navigation most studies focus on representations of one-step relationships between visited locations.

Indeed if mammalian path integration was merely computed online using a map of one-step transitions, one would not expect firing distance-to-goal cells, like the goal-vector cells shown in bats (Sarel et al., 2017b), reward cells shown in rodents (Gauthier & Tank, 2018), or or trace cells in humans representing distances to remembered locations (Qasim et al., 2018). However, recapturing the idea of cognitive maps in terms of predictive representations of multi-step relationships can offer a different view on the relationship between map-based and vector-based navigation. The correspondence of our simulated distance-to-goal firing and the finding of vectorial representations in the bat suggests that a linear function over multi-scale predictive maps is - at least partly - consistent with a vector-based representation of goals (Kubie & Fenton, 2009; Bush et al., 2015). Intuitively, this makes sense in light of previous computational proposals for vectorial representations.

It has been proposed that in order to accomplish vector-based navigation the brain computes and stores a “shortcut matrix” using path integration (Kubie & Fenton, 2009), leading to egocentric and immediate firings of direction and distance to goal without further online computations. While a shortcut matrix does not readily arise from traditional one-step cognitive maps since Tolman (Tolman, 1948), the sorts of representations computed for a shortcut matrix are already learned in the multi-step dependencies that are stored in the successor representation (Dayan, 1993; Momennejad, Russek, et al., 2017). First, consistent with the idea of a shortcut matrix, relationships between states that are multiple steps apart are cached, and second, as we show here, predictions consistent with the firing of cells in specific distances to goal states can be derived from the multi-scale SR account. Our proposal could potentially extend the shortcut vector view to multi-scale or hierarchical shortcut vectors, where rows of SRs at different scales correspond to shortcut vectors at hierarchically different levels of abstraction – well-suited for complex navigation. Future work is required to systematically compare the mathematical relationship between the two accounts, as well as how they compare when simulating empirical data.

Our proposal is also consistent with recent work by Banino, Barry and colleagues, who have shown that training a recurrent network to perform path integration leads to the emergence of grid-like representations (Banino et al., 2018). Importantly, deep reinforcement learning agents using such grid-like representation and vectorial navigation outperformed other models (and humans) in goal-directed navigation in unfamiliar environments, and mammal-like discovery of shortcuts. We showed that the distance to goal aspect of vectorial representations appears readily using the derivative of multi-scale SR, through the activation of cells corresponding to different neighboring horizons of the destination state (corresponding to different σ values). The dependence on head direction can perhaps be captured in the policy-dependent aspect of successor representations as well the relationship to the grid system and basis set/eigen-vector accounts (Gustafson & Daw, 2011; Stachenfeld et al., 2017). However, though every SR matrix is computed under a specific policy toward the goal state, here we have assumed SRs with a single policy for simplification. The direction to goal aspect of the vector-based representation is, thus, less readily apparent in the present version of multi-scale SR. Future accounts can expand this model to systematically simulate policy-dependent SRs and their relationship to simulating head-direction cells during navigation.

### Future directions

#### Policy dependence

One of the main features of SR is that it is policy dependent. That is, SR matrices are computed given particular action policies that the agent takes, rather than the alocentric map of the environment. SR is not optimal for facilitating “full model-based,” off-policy, alocentric, action-outcome contingent analyses. Due to policy-dependence, SR over-represents parts of the state space that are often visited under its corresponding policy. Perhaps a rough analogy is that of walking in the snow: paths that are more often taken are carved deeper and more clearly than paths less traveled. This analogy can explain why the direction to often-visited states (or goal states with a high value) should readily arise from the neural representation of multi-scale SRs. That said, in principle it is possible that SRs at different time-scales are formed with different policies as well, corresponding to the statistics of the environment at different time-scales. If true, this could allow flexible policy-choice depending on environmental statistics and the temporal horizon of the problem at hand. That is, policy-dependence could potentially lead to selecting different policy depending on the time-scale or planning horizon.

One possibility is that different policies mark the relevant SR at different scales of abstraction, and at the moment of decision making, arbitration between representations from different policies might be required. In the case of navigation, the policy-dependence of SRs, and potentially different policies used for abstraction at different levels, may yield interesting relationships to head-direction cells and grid cells when choosing policy at different scales or horizons for planning future actions. Future studies can be designed to model and test specific predictions of a multi-scale policy dependent model.

#### Expanding the SR ensemble via offline replay

To overcome limited predictive and planning horizon, we have proposed that the brain simultaneously caches multiple successor representations with different discount rates (Figure 1). However, attentional and learning resources may be limited to specific horizons during experience and online learning. As one increases the number of scales (γs) at which SR matrices are computed, the resolution for the estimate over future time points increases. This resolution comes at a high cost: the amount of resources committed to constructing the representation ‘online’ goes up proportional to the number of scales. This means that, given a fully online strategy, the system must commit the resources necessary to construct the multi-scale representations at the outset of learning. However, in the absence of an *a priori* expectation about the meta-parameters of the environment or information about what scale—or scales—will be relevant in a particular learning problem, the cost of learning representations at multiple scales may be too high.

To overcome this further limitation, one possibility is that offline replay enables the brain to learn and cache more successor representations with a larger set of discount rates ‘offline’. Depending on the parameters of the task, this could potentially enable better separation of different categories, better integration and inference (Momennejad, Otto, Daw, & Norman, 2017), or clustering and generalization of items from the same context. We have elsewhere proposed and named the combination of offline replay and the successor representation SR-Dyna (Momennejad, Russek, et al., 2017; Russek et al., 2017). A multi-scale extension of SR-Dyna could use offline replay to cache representations at scales beyond what has been learned during direct experience. This further proposal for a multi-scale SR-Dyna remains to be computationally fleshed out and empirically tested in future studies.

## Summary

We propose a multi-scale model of predictive representations that overcomes caveats in existing models incorporating the successor representation. Computing the order and sequence of future states following a starting state is non-trivial and costly: it requires nonlinear operations on entire SR matrices that can be large. Here we show that the derivatives of multiple successor representations (computed with different scales of abstraction) can be used to recover distance and sequential order of successor states, reconstructing entire expected future trajectories. Our proposal offers a mechanism for constructing an estimated timeline of sequential future events using representations abstracted at multiple time-scales. This proposal expands previous work on predictive representations in reinforcement learning (Momennejad, Russek, et al., 2017; Russek et al., 2017; Stachenfeld et al., 2017) and has the same properties as an analogous representation of remembering or reconstructing a timeline for the past (Shankar & Howard, 2013). The model’s prediction of distance-to-goal cells is in line with recent findings in the bat, rodent, and human literature (Sarel et al., 2017b; Gauthier & Tank, 2018; Qasim et al., 2018), and may offer promising insight into a unified model of cognitive map and vector-based representations (Kubie & Fenton, 2009; Banino et al., 2018) used in navigation and planning. This proposal can be applied to planning in the context of spatial navigation as well as non-spatial cognition. We expect the model to inspire future theoretical and empirical studies in reinforcement learning, navigation, and planning.

## Acknowledgments

We thank Sam Gershman for influential early discussions and also thoughtful reactions to an earlier draft of the manuscript. We especially thank Kim Stachenfeld for helpful conversations and Per Sederberg, Zoran Tiganj, Salman Qasim, Nathaniel Daw, and Dylan Rich for useful discussions. This work was funded by the John Templeton Foundation (IM),NIBIB R01EB022864, and ONR MURI N00014-16-1-2832 (MWH).

## References

Akhlaghpour, H., Wiskerke, J., Choi, J. Y., Taliaferro, J. P., Au, J., & Witten, I. (2016). Dissociated sequential activity and stimulus encoding in the dorsomedial striatum during spatial working memory. eLife, 5, e19507.

Aronov, D., Nevers, R., & Tank, D. W. (2017, March). Mapping of a non-spatial dimension by the hippocampalentorhinal circuit. Nature, 543 (7647), 719. Retrieved 2018-05-29, from https://www.nature.com/articles/nature21692 doi: doi:10.1038/nature21692

Banino, A., Barry, C. J., Benigno, U., Blundell, C., Lillicrap, T., Mirowski, P., … Kumaran, D. (2018, May). Vector-based navigation using grid-like representations in artificial agents. Nature. Retrieved 2018-08-27, from dx.doi.org/10.1038/s41586-018-01026

Barry, C., Hayman, R., Burgess, N., & Jeffery, K. J. (2007). Experience-dependent rescaling of entorhinal grids. Nature Neuroscience.

Bell, A. J., & Sejnowski, T. J. (1997). The independent components of natural scenes are edge filters. Vision research, 37(23), 3327–3338.

Botvinick, M., & Weinstein, A. (2014, November). Model-based hierarchical reinforcement learning and human action control. Philos. Trans. R. Soc. Lond., B, Biol. Sci., 369 (1655). doi: 10.1098/rstb.2013.0480

Brun, V. H., Solstad, T., Kjelstrup, K. B., Fyhn, M., Witter, M. P., Moser, E. I., & Moser, M.-B. (2008). Progressive increase in grid scale from dorsal to ventral medial entorhinal cortex. Hippocampus, 18(12), 1200–1212. doi: 10.1002/hipo.20504

Brunec, I. K., Bellana, B., Ozubko, J. D., Man, V., Robin, J., Liu, Z.-X., … Moscovitch, M. (2018). Multiple scales of representation along the hippocampal anteroposterior axis in humans. Current Biology, 28(13), 2129–2135.e6. doi: 10.1016/j.cub.2018.05.016

Bush, D., Barry, C., Manson, D., & Burgess, N. (2015, August). Using Grid Cells for Navigation. Neuron, 87(3), 507–520. Retrieved 2018-08-25, from www.cell.com/neuron/abstract/S0896-6273(15)00628-5 doi: 10.1016/j.neuron.2015.07.006

Collin, S. H., Milivojevic, B., & Doeller, C. F. (2015). Memory hierarchies map onto the hippocampal long axis in humans. Nature Neuroscience.

Dayan, P. (1993, July). Improving Generalization for Temporal Difference Learning: The Successor Representation. Neural Computation, 5(4), 613–624. Retrieved from https://doi.org/10.1162/neco.1993.5.4.613 doi: 10.1162/neco.1993.5.4.613

Dayan, P., & Balleine, B. W. (2002, October). Reward, Motivation, and Reinforcement Learning. Neuron, 36(2), 285–298. Retrieved 2018-09-18, from http://www.sciencedirect.com/science/article/pii/S0896627302009637 doi: 10.1016/S0896-6273(02)00963-7

Eichenbaum, H., Kuperstein, M., Fagan, A., & Nagode, J. (1987, March). Cue-sampling and goal-approach correlates of hippocampal unit activity in rats performing an odor-discrimination task. J. Neurosci., 7(3), 716–732. Retrieved 2018-09–15, from http://www.jneurosci.org/content/7/3/716 doi: 10.1523/JNEUROSCI.07-0300716.1987

Gardner, M., Schoenbaum, G., & Gershman, S. J. (2018). Rethinking dopamine prediction errors. bioRxiv, 239731.

Garvert, M. M., Dolan, R. J., & Behrens, T. E. (2017). A map of abstract relational knowledge in the human hippocampal–entorhinal cortex. eLife, 6, e17086.

Gauthier, J. L., & Tank, D. W. (2018). A dedicated population for reward coding in the hippocampus. Neuron.

Gershman, S. J. (2018). The successor representation: its computational logic and neural substrates. Journal of Neuroscience, 0151–18.

Gershman, S. J., Moore, C. D., Todd, M. T., Norman, K. A., & Sederberg, P. B. (2012, June). The Successor Representation and Temporal Context. Neural Computation, 24(6), 1553–1568.

Gustafson, N. J., & Daw, N. D. (2011, October). Grid cells, place cells, and geodesic generalization for spatial reinforcement learning. PLoS Comput. Biol., 7(10), e1002235. doi: 10.1371/journal.pcbi.1002235

Howard, M. W., MacDonald, C. J., Tiganj, Z., Shankar, K. H., Du, Q., Hasselmo, M. E., & Eichenbaum, H. (2014, March). A Unified Mathematical Framework for Coding Time, Space, and Sequences in the Hippocampal Region. J Neurosci, 34(13), 4692–4707. Retrieved from https://www.ncbi.nlm.nih.gov/pmc/articles/PMC3965792/ doi: 10.1523/JNEUROSCI.5808-12.2014

Jin, D. Z., Fujii, N., & Graybiel, A. M. (2009). Neural representation of time in cortico-basal ganglia circuits. Proceedings of the National Academy of Sciences, 106(45), 19156–19161.

Johnson, A., & Redish, A. D. (2007). Neural ensembles in CA3 transiently encode paths forward of the animal at a decision point. Journal of Neuroscience, 27(45), 12176–89.

Jung, M. W., Wiener, S. I., & McNaughton, B. L. (1994). Comparison of spatial firing characteristics of units in dorsal and ventral hippocampus of the rat. Journal of Neuroscience, 14(12), 7347–56.

Kjelstrup, K. B., Solstad, T., Brun, V. H., Hafting, T., Leutgeb, S., Witter, M. P., … Moser, M. B. (2008a). Finite scale of spatial representation in the hippocampus. Science, 321(5885), 140–3.

Kjelstrup, K. B., Solstad, T., Brun, V. H., Hafting, T., Leutgeb, S., Witter, M. P., … Moser, M.-B. (2008b, July). Finite scale of spatial representation in the hippocampus. Science, 321(5885), 140–143. doi: 10.1126/science.1157086

Kraus, B. J., Brandon, M. P., Robinson, R. J., Connerney, M. A., Hasselmo, M. E., & Eichenbaum, H. (2015). During running in place, grid cells integrate elapsed time and distance run. Neuron, 88 (3), 578–589.

Kubie, J. L., & Fenton, A. A. (2009, May). Heading-vector navigation based on head-direction cells and path integration. Hippocampus, 19(5), 456–479. Retrieved 2018-08-27, from https://www.onlinelibrary.wiley.com/doi/abs/10.1002/hipo.20532 doi: 10.1002/hipo.20532

Kurth-Nelson, Z., & Redish, A. D. (2009). Temporal-difference reinforcement learning with distributed representations. PLoS One, 4(10), e7362.

Liu, Y., Tiganj, Z., Hasselmo, M. E., & Howard, M. W. (in press). Biological simulation of scale-invariant time cells biological simulation of scale-invariant time cells. Hippocampus.

Lubenov, E. V., & Siapas, A. G. (2009). Hippocampal theta oscillations are travelling waves. Nature, 459 (7246), 534–9.

Ludvig, E. A., Sutton, R. S., & Kehoe, E. J. (2008). Stimulus representation and the timing of reward-prediction errors in models of dopamine system. Neural Computation, 20, 3034–3054.

MacDonald, C. J., Lepage, K. Q., Eden, U. T., & Eichenbaum, H. (2011). Hippocampal “time cells” bridge the gap in memory for discontiguous events. Neuron, 71(4), 737–749.

Mao, D., Neumann, A. R., Sun, J., Bonin, V., Mohajerani, M. H., & McNaughton, B. L. (2018, July). Hippocampus-dependent emergence of spatial sequence coding in retrosplenial cortex. PNAS, 201803224. Retrieved 2018-10-05, from http://www.pnas.org/content/early/2018/07/13/1803224115 doi: 10.1073/pnas.1803224115

Marr, D., & Hildreth, E. (1980). Theory of edge detection. Proceedings of the Royal Society of London B, 207(1167), 187–217.

Mau, W., Sullivan, D. W., Kinsky, N. R., Hasselmo, M. E., Howard, M. W., & Eichenbaum, H. (2018, May). The Same Hippocampal CA1 Population Simultaneously Codes Temporal Information over Multiple Timescales. Current Biology, 28(10), 1499–1508.e4. Retrieved 2018-10-22, from http://www.sciencedirect.com/science/article/pii/S0960982218303804 doi: 10.1016/j.cub.2018.03.051

Meister, M. L., & Buffalo, E. A. (2017). Conjunctive coding in the primate entorhinal cortex. In Society for neuroscience abstracts (Vol. 425.23).

Mello, G. B., Soares, S., & Paton, J. J. (2015). A scalable population code for time in the striatum. Current Biology, 25(9), 1113–1122.

Momennejad, I., Otto, A. R., Daw, N. D., & Norman, K. A. (2017). Offline Replay Supports Planning: fMRI Evidence from Reward Revaluation. bioRxiv. Retrieved from https://www.biorxiv.org/content/early/2017/10/02/196758 doi: 10.1101/196758

Momennejad, I., Russek, E. M., Cheong, J. H., Botvinick, M. M., Daw, N. D., & Gershman, S. J. (2017, September). The successor representation in human reinforcement learning. Nature Human Behaviour, 1(9), 680–692. Retrieved 2018-05-29, from https://www.nature.com/articles/s41562-017-0180-8 doi: 10.1038/s41562-017-0180-8

Olshausen, B. A., & Field, D. J. (1996). Emergence of simple-cell receptive field properties by learning a sparse code for natural images. Nature, 381 (6583), 607–609.

Omer, D. B., Maimon, S. R., Las, L., & Ulanovsky, N. (2018, January). Social place-cells in the bat hippocampus. Science, 359 (6372), 218–224. Retrieved 2018-10-02, from http://science.sciencemag.org/content/359/6372/218 doi: 10.1126/science.aao3474

Pastalkova, E., Itskov, V., Amarasingham, A., & Buzsaki, G. (2008). Internally generated cell assembly sequences in the rat hippocampus. Science, 321 (5894), 1322–7.

Patel, J., Fujisawa, S., Berényi, A., Royer, S., & Buzsáki, G. (2012). Traveling theta waves along the entire septotemporal axis of the hippocampus. Neuron, 75(3), 410–7. doi: 10.1016/j.neuron.2012.07.015

Pfeiffer, B. E., & Foster, D. J. (2013). Hippocampal place-cell sequences depict future paths to remembered goals. Nature, 497 (7447), 74–9. doi: 10.1038/nature12112

Post, E. (1930). Generalized differentiation. Transactions of the American Mathematical Society, 32, 723–781.

Qasim, S. E., Miller, J., Inman, C. S., Gross, R., Willie, J. T., Lega, B., … Jacobs, J. (2018). Neurons remap to represent memories in the human entorhinal cortex. bioRxiv. Retrieved from https://www.biorxiv.org/content/early/2018/10/16/433862 doi: 10.1101/433862

Russek, E. M., Momennejad, I., Botvinick, M. M., Gershman, S. J., & Daw, N. D. (2017, September). Predictive representations can link model-based reinforcement learning to model-free mechanisms. PLOS Computational Biology, 13(9), e1005768. Retrieved 2018-05-29, from http://journals.plos.org/ploscompbiol/article?id=10.1371/journal.pcbi.1005768 doi: 10.1371/journal.pcbi.1005768

Sarel, A., Finkelstein, A., Las, L., & Ulanovsky, N. (2017a). Vectorial representation of spatial goals in the hippocampus of bats. Science, 355(6321), 176–180.

Sarel, A., Finkelstein, A., Las, L., & Ulanovsky, N. (2017b, January). Vectorial representation of spatial goals in the hippocampus of bats. Science, 355(6321), 176–180. Retrieved 2018-05-29, from http://science.sciencemag.org/content/355/6321/176 doi: 10.1126/science.aak9589

Sarel, A., Finkelstein, A., Las, L., & Ulanovsky, N. (2017c, January). Vectorial representation of spatial goals in the hippocampus of bats. Science, 355(6321), 176–180. Retrieved 2018-07-03, from http://science.sciencemag.org/content/355/6321/176 doi: 10.1126/science.aak9589

Schapiro, A. C., Rogers, T. T., Cordova, N. I., Turk-Browne, N. B., & Botvinick, M. M. (2013, April). Neural representations of events arise from temporal community structure. Nature Neuroscience, 16(4), 486–492. Retrieved 2018-08-28, from https://www.nature.com/articles/nn.3331 doi: 10.1038/nn.3331

Shankar, K. H., & Howard, M. W. (2012). A scale-invariant internal representation of time. Neural Computation, 24(1), 134–193.

Shankar, K. H., & Howard, M. W. (2013). Optimally fuzzy temporal memory. Journal of Machine Learning Research, 14, 3753–3780.

Shankar, K. H., Singh, I., & Howard, M. W. (2016). Neural mechanism to simulate a scale-invariant future. Neural Computation, 28, 2594–2627.

Smith, T. A., Hasinski, A. E., & Sederberg, P. B. (2013). The context repetition effect: Predicted events are remembered better, even when they dont happen. Journal of Experimental Psychology: General, 142, 1298–1308.

Spears, T. A., Jacques, B. G., Howard, M. W., & Sederberg, P. B. (2017). Scale-invariant temporal history (sith): optimal slicing of the past in an uncertain world. arXiv preprint arXiv:1712.07165.

Stachenfeld, K. L., Botvinick, M. M., & Gershman, S. J. (2017, November). The hippocampus as a predictive map. Nature Neuroscience, 20(11), 1643–1653. Retrieved 2018-05-29, from https://www.nature.com/articles/nn.4650 doi: 10.1038/nn.4650

Tiganj, Z., Cromer, J. A., Roy, J. E., Miller, E. K., & Howard, M. W. (2018). Compressed timeline of recent experience in monkey lPFC. Journal of Cognitive Neuroscience, 30, 935–950.

Tiganj, Z., Gershman, S. J., Sederberg, P. B., & Howard, M. (2018). Estimating scale-invariant future in continuous time. arXiv, arXiv:1802.06426.

Tiganj, Z., Kim, J., Jung, M. W., & Howard, M. W. (2017). Sequential firing codes for time in rodent mPFC. Cerebral Cortex, 27, 5663–5671.

Tolman, E. C. (1948, Jul). Cognitive maps in rats and men. Psychological Review, 55(4), 189–208.

Tsao, A., Sugar, J., Lu, L., Wang, C., Knierim, J. J., Moser, M.-B., & Moser, E. I. (2017). Integrating time in lateral entorhinal cortex. In Society for neuroscience abstracts (Vol. 84.21).

Tsao, A., Sugar, J., Lu, L., Wang, C., Knierim, J. J., Moser, M.-B., & Moser, E. I. (2018, August). Integrating time from experience in the lateral entorhinal cortex. Nature, 1. Retrieved 2018-08-29, from https://www.nature.com/articles/s41586-018-0459-6 doi: 10.1038/s41586-018-0459-6

Zhang, H., & Jacobs, J. (2015, Sep). Traveling theta waves in the human hippocampus. Journal of Neuroscience, 35(36), 12477–87. doi: 10.1523/JNEUROSCI.5102-14.2015

